# The impact of fire intensity on plant growth forms in high-altitude Andean grassland

**DOI:** 10.1101/2020.04.25.061051

**Authors:** Maya A. Zomer, Paul M. Ramsay

## Abstract

Fires in the páramo grasslands of the tropical northern Andes vary in intensity at the landscape scale. Fire suppression strategies, intended to conserve biodiversity and páramo ecosystem integrity and function, could lead to the accumulation of high fuel loads and ultimately fires of higher intensity. Yet the impact of fire intensity on páramos is not well studied or understood. 5½ years after a fire, we measured plant growth form composition, light transmission to the ground and soil temperature in plots representing very high, high, medium, and low fire intensities, plus a “control” that had been unburned for at least 40 years. We also assessed *Espeletia* rosette diameters, heights, population density, and mortality. The low intensity plot, with a closed canopy of vegetation and lower growth form diversity, contrasted with the very high intensity plot, with patchy vegetation cover and higher growth form diversity. The high intensity plot had shorter *Espeletia* plants with smaller rosettes. Light transmission to the ground increased with fire intensity, as did soil temperatures. We demonstrate that the same fire can produce different microenvironmental conditions, plant communities, and population structures in different parts of the same fire event. In future, fire suppression could provoke more intense fires with consequences for ecosystem function and service provision. Since intensity is determined by a complex interaction of factors, we advocate a field programme of experimental fires for a better understanding of páramo fire ecology and to guide effective páramo conservation strategies.

## Introduction

The high-altitude grasslands of the northern Andes, known as páramos, represent the largest extension of tropical alpine ecosystems, extending between the upper treeline and the perennial snowline (about 3200–5000 m altitude) and covering an area of around 35,000 km^2^ (Madriñán et al. 2013). These biodiverse tropical alpine regions provide important ecological services such as carbon storage and water supply for people and ecosystems at lower elevations. (Buytaert et al. 2011; Ramsay 2014).

Fires have been present in the Andean páramos for thousands of years and are of fundamental significance to the biodiversity patterns and ecological processes of these grasslands (Horn and Kappelle 2009; White 2013). Modern páramo fires are predominantly set by local farmers to encourage new growth for the grazing of livestock. The frequency of burning depends on vegetation recovery but is typically every 2–5 years (Ramsay and Oxley 1996). This has resulted in a mosaic pattern of fire patches at the landscape scale (Laegaard 1992).

Land use in the páramos has shifted in the last two decades, with an intensification of agricultural and recreational use, putting significant pressure on these ecologically important, high diversity systems (Armenteras et al. 2020; Ramsay 2014). As a response, some páramo regions have adopted a fire suppression management strategy to conserve páramo ecosystem integrity and function. A possible unintended consequence of suppression of fires in grassland landscapes is the accumulation of high fuel loads (Keating 2007). Fuel load is the total amount of dry biomass or “the total amount of heat energy available for release during fires” (Whelan 1995). This could potentially lead to less frequent, widespread fires of high intensity that might have significant effects on the landscape (Keating 2007). An understanding of fire intensity and ecosystem response to changes in páramo fire regimes is therefore essential to inform current conservation efforts and management decisions.

The behaviour of páramo fires is highly variable and is determined by factors such as fuel availability, wind direction, site topography, vegetation composition and structure, climatic conditions and human intervention (Horn and Kappelle 2009). Fire intensity refers to energy output during various stages of the fire, including several characteristics such as temperature, residence time, radiant temperature, flame length, and fireline intensity (Keeley 2009; Rossi et al. 2018). Fireline intensity is the most commonly used metric of fire intensity and is defined by Byram (1959) as the rate of heat release per unit time per unit length of fire front. Fire severity refers to the post fire loss or change in organic matter above and below ground (Díaz-Delgado et al. 2003; Keeley 2009). Plant mortality is one metric of fire severity, but only if the mortality is the genuine death of the plant rather than temporary loss of material (Keeley 2009). Thus, fire intensity and severity are distinct from one another, but correlated. The most basic understanding of this is that the longer plants are exposed to high temperatures, the more damaging and severe the effects of fire (Ghermandi et al. 2013; Whelan 1995).

Ramsay and Oxley (1996) measured temperatures of a páramo fire and found that temperatures varied depending on the position within the vegetation structure. Maximum temperatures were highest in the upper leaves of tussock grasses (sometimes >500 °C), but were low (<65°C) amongst the dense leaf bases and just below ground. Thus, plant survival is actually not only dependent on the temperature and duration of fire, but also on the position of a plant’s crucial tissues within this tussock structure (Ramsay and Oxley 1996). Post-fire changes in organic matter and post-fire plant community composition therefore reflects fire event conditions combined with the differential survival, recovery and recruitment of plant species (Keeley and Fotheringham 2000). As a result, post-fire ecosystem responses are highly variable in the páramos (Keating 2007; Luteyn 1999; Sklenář and Ramsay 2001; Suárez and Medina 2001). While the roles of fire intensity and severity are widely acknowledged to be important factors in these varying outcomes, the topic is not well studied or understood in páramo ecosystems (Bremer et al. 2019; Horn and Kappelle 2009; Ramsay and Oxley 1996).

The grass páramo of northern Ecuador and Colombia is dominated by tussock grasses and giant rosettes of *Espeletia* (Asteraceae). *Espeletia* or ‘frailejon’ is a flagship plant of the páramos (even represented for several years on the Colombian 100 peso coin) and merits particular attention for this reason. Laegaard (1992) suggested that *Espeletia* germination and establishment are enhanced after fire opens up the canopy and allows more light to reach the ground. Mature *Espeletia* plants are resilient to fire because their stems retain dead leaves which provide an insulating layer against lethal fire temperatures, and which are packed so tightly that there is not enough oxygen to sustain burning. The dead leaf cover on *Espeletia* stems has been reported as a reliable indicator of time since fire in the páramos (Zomer and Ramsay 2018). However, repeated fires, each removing a little more of the dead leaves, expose the stem below. Once the layer of dead leaf bases is destroyed, the plant is thought to be more vulnerable to fire (Laegaard 1992; Smith 1981). Taller (older) plants would normally have been subjected to more fires in their lifetime than shorter plants, and are therefore more susceptible to mortality (Ramsay 2014; Smith 1981).

A rich flora accompanies the dominant tussocks and rosettes, and this diversity makes it difficult to study and interpret community-level successional trends at species level. Many páramo plants have protected buds, rhizomes or rootstock that are able to regenerate after fire even when the rest of the plant has been destroyed, but the mechanisms vary from species to species (Laegaard 1992). Grouping species into functional types that are similar in their resource use and responses to environmental controls (Wilson 1999) has proved useful in other circumstances (Duckworth et al. 2000; Smith and Huston 1989). With broad overview studies in mind, Ramsay and Oxley (1997) proposed ten plant growth forms for the páramos: stem rosettes, basal rosettes, tussock grasses, acaulescent rosettes, cushions and mats, upright shrubs, prostrate shrubs, erect herbs, prostrate herbs and trailing herbs.

Our study takes advantage of observations made by Ramsay (2014) during a single páramo fire event in northern Ecuador, and putative fire intensities of very high, high, medium, and low intensities were assigned to plots all within 600 m of one another. The fire was extinguished manually and an area at the point where the fire was stopped offered a “control” which had been unburned for at least 40 years. We revisited these same plots 5½ years after the fire. We aimed to determine the impact of fire intensity on the vegetation and related microclimatic conditions (light availability and temperature). We also considered the impact of fire intensity on the relative abundance of plant growth forms, with particular interest in *Espeletia* because of its flagship status.

## Methods

### 2.1 Study areas

The Reserva Ecológica El Ángel (REEA), in northern Ecuador, contains approximately 16,000 ha of páramo covering altitudes ranging from 3400m to 4200m. The grass páramos in the reserve are dominated by tussocks of *Calamagrostis intermedia* (J.Presl) Steud. and giant rosettes of *Espeletia pycnophylla* Cuatrec. This area has experienced direct impacts from human activities, including cattle grazing and regular burning of the páramo, as well as conversion of land for cultivation (Moscol Olivera and Cleef 2009). Although the reserve has banned fires since it was established in 1992, fires are still common in certain parts of the buffer zone and adjoining areas within the reserve itself.

Ramsay (2014) had the opportunity to observe a single fire that burned through the grass páramo at 3600 m in the buffer zone of REEA on 3 August 2009. As the fire burned, qualitative observations were made, and putative fire intensities were assigned, representing a relative gradient of very high, high, medium, and low intensities (Ramsay 2014). These fire intensity areas burned at approximately the same time (within 2–3 hours) and were all located within 600 m of one another. The fireline was manually extinguished near the high and very high intensity plots, preventing nearby vegetation from burning. Within 10 m of this fire limit, a plot was established to represent an unburned control. Based on the dead leaf cover of *Espeletia* plants within this plot (see Zomer and Ramsay 2018), this plot had not been burned for at least 40 years, and was very similar in structure and composition before the fire to the neighbouring high and very high intensity plots. Ramsay (2014) observed the immediate effects of different fire intensities on *Espeletia* populations and found that the numbers of standing dead plants was clearly correlated with the assigned fire intensities.

### 2.2 Field Measurements and Data Analysis

GPS coordinates were used to locate precisely five areas identified by Ramsay (2014) 5½ years after the August 2009 fire. Representative photographs of these areas, taken during our survey work, are shown in Fig. 1 (which can be compared with the photographs in Fig. 1 of Garcia-Meneses and Ramsay 2014, showing the plots immediately after the fire).

**Fig. 1.**
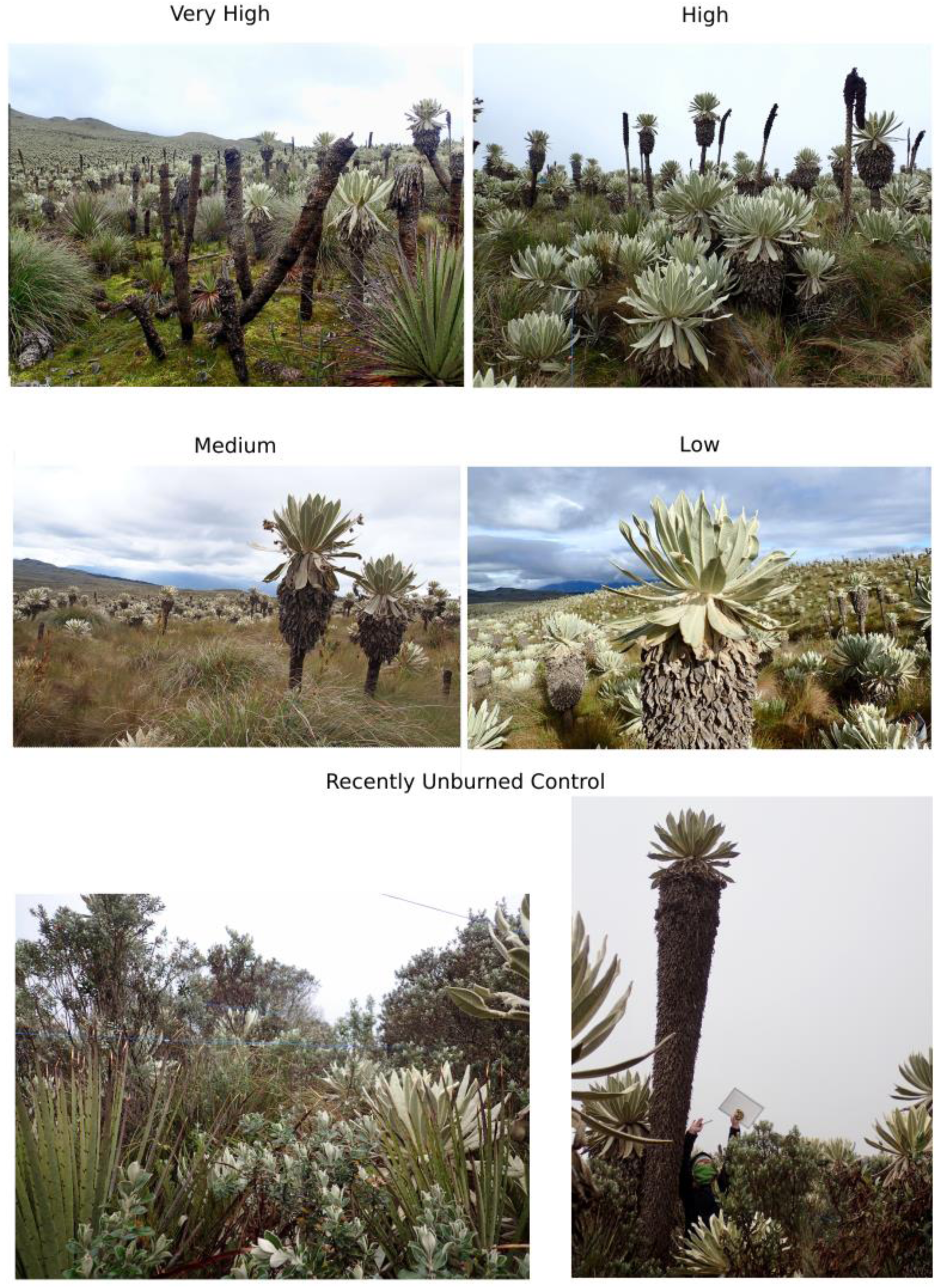
Plots of different fire intensity in April 2015, 5½ years after the August 2009 fire. Photos: Maya Zomer.

A 50 m x 2 m plot was established within each area. The plot was divided into one hundred 1 m^2^ quadrats. The presence of ten páramo plant growth forms (Ramsay and Oxley 1997) was recorded for each 1 m^2^ quadrat. The number of *Espeletia* giant rosette plants was counted within each plot, along with each plant’s height, rosette diameter and whether it was dead or alive. Plants were only included if more than half of the stem, where it entered the ground, was located within the plot.

Vegetation height (excluding *Espeletia* plants) was measured in each 1 m^2^ quadrat using the drop-disc method (Stewart et al. 2001), with the resting height of a 20 cm diameter disc, 190 g in weight, taken as the vegetation height.

Light reaching the ground was determined by comparing measurements of incident photosynthetically active radiation (PAR) using a SunScan Canopy Analysis System with BF2 Beam Fraction Sensor (Delta-T Devices Ltd, Cambridge, UK). The SunScan probe was held at ground level underneath the vegetation at fifty 1 m intervals along the length of the plot. Each reading consisted of 64 separate light measurements of the percentage of total light above the canopy reaching the sensors on the ground. The mean percentage of total light reaching ground level at each 1 m interval was calculated, and subsequent analyses were based on these mean values (*n*=50 for each plot).

Soil temperature was measured at 20 cm soil depth using Signstek 3 1/2 6802 II Dual Channel Digital Thermometer with 2 k Type Thermocouple Sensor Probes. Measurements were taken at five regular intervals along the longest axis of the plot, at 5 m, 15 m, 25 m, 35 m, and 45 m. At each interval, a temperature reading was taken in three positions of different shading levels: beneath dense tussocks (closed canopy), on the edge of tussocks (intermediate) and in open intertussock areas (open canopy).

The Shannon Diversity Index (using log_e_) and the Gini-Simpson Diversity Index (which particularly emphasizes the role of dominance in diversity) were calculated for growth forms in all plots, using Primer 6 (PRIMER-E, Plymouth, UK). We also used non-metric multidimensional scaling (MDS) in Primer 6 to compare square-root-transformed abundances of growth forms between the fire intensity plots. Other standard statistical tests were performed with R (R Core Team 2019).

## Results

Vegetation height (excluding *Espeletia*) was much taller, and more variable, in the control plot than in the burned ones, while the very high intensity plot had the shortest vegetation (Kruskal-Wallis Test, *X*_4_^2^ = 100.21, *p* < 0.0001; Fig. 2A). The vegetation in the control plot was taller than 2 m in more than one-quarter of the measurements, while all measurements in the burned plots were less than 1.2 m. More than half the measurements of vegetation height in the very intense fire plot were less than 12.5 cm.

**Fig. 2.**
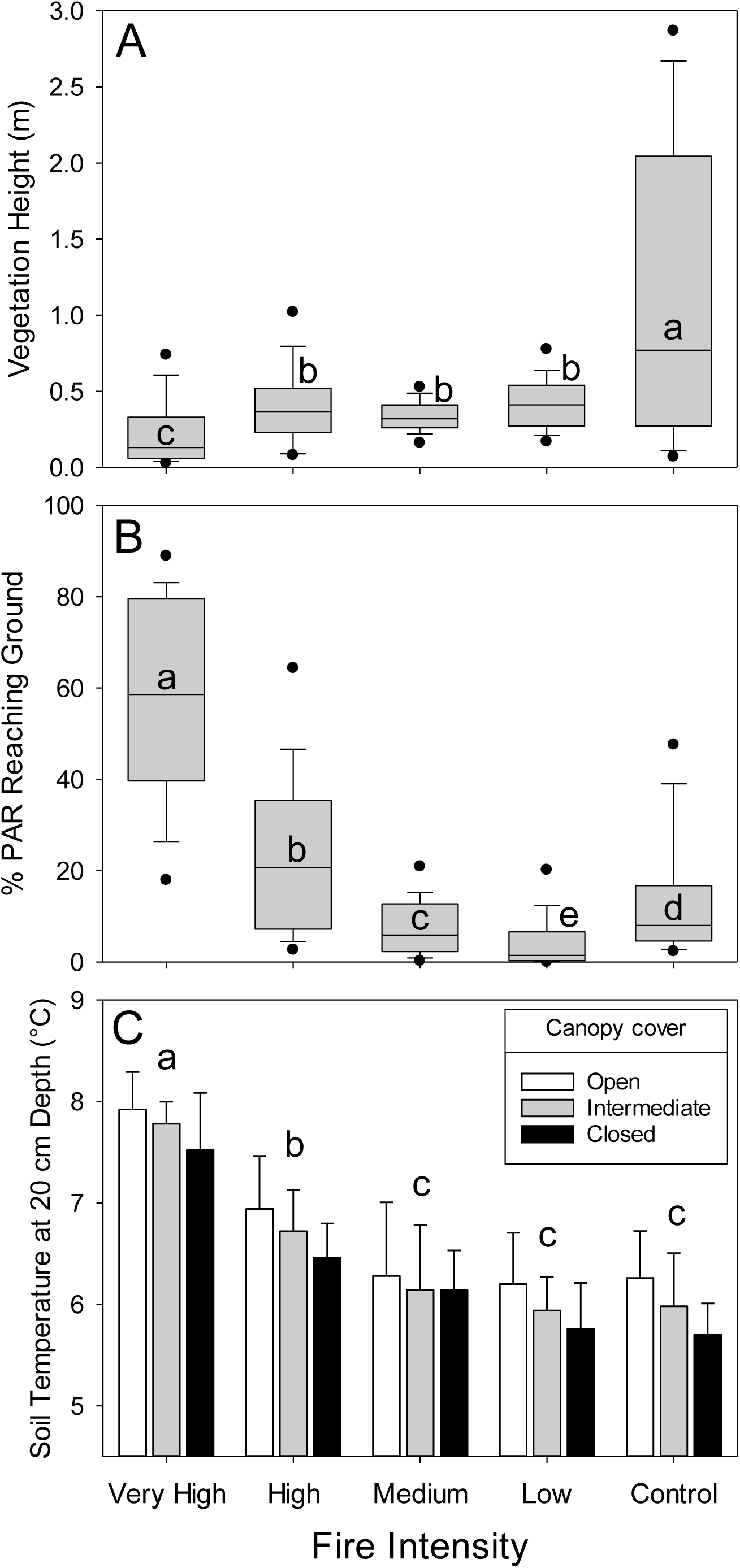
(A) Vegetation height (excluding *Espeletia, n*=100 in each case), (B) light reaching the ground (*n*=50 in each case), and (C) soil temperatures at 20 cm depth (*n*=15 in each case) in plots of different fire intensities, 5½ years after the fire. Medians are shown as central lines, except in (C) which shows means, boxes are 25^−^75^th^ percentiles, whiskers are 10–90^th^ percentiles, and the dots show 5–95^th^ percentiles. Medians/means sharing a letter in each panel were not significantly different (*p*≤0.05).

We recorded most light transmission through the canopy to the ground in the very high intensity plot, while the least proportion of available light reached the ground in the low intensity fire plot, with the others intermediate (Kruskal-Wallis Test *X*_4_^2^ = 143.50, *p* < 0.0001; Fig. 2B). The median measure of PAR transmission measurement through the vegetation canopy was 60% in the very high intensity fire plot, considerably higher than any other plot. While none of the PAR transmission measurements in the very high intensity fire plot fell below 10%, the majority of the measurements in the low intensity fire plot had light transmission at or below that value.

Soil temperature at 20 cm depth was, on average, 0.4 °C higher in places with an open canopy compared with places with a closed canopy, while intermediate canopy cover were not significantly different from open or closed canopies (GLM ANOVA, Canopy *F*_2,60_ = 4.641, *p* < 0.0134; Fig.5C). Soil temperatures were highest in the very high intensity fire plot, with the medium intensity, low intensity and unburned control plots similarly low, while the high intensity plot was intermediate (GLM ANOVA, Intensity *F*_4,60_ = 38.069, *p* < 0.0001; Fig. 2C). The difference in temperature between these two groups was approximately 1.75 °C. There was no interaction of soil temperature between fire intensity and vegetation canopy cover (GLM ANOVA, Intensity x Canopy *F*_8,60_ = 0.036, *p* = 0.9949).

Plant growth form composition differed among the fire intensity plots (Table 1). Tussocks were almost ubiquitous in all but the very high intensity and unburned control plots, where they were nevertheless very abundant. Giant stem rosettes were also very abundant in all plots. Upright shrubs, cushions and mats, prostrate herbs, and giant basal rosettes were most abundant in the very high intensity plot, and least abundant in the low intensity plot. Erect herbs and prostrate shrubs showed a similar pattern, though their highest abundances were in the medium intensity plots. Acaulescent rosettes were found in low abundance in all burned plots, and were absent from the unburned control, while trailing herbs were absent from all burned plots, and were only found, in low abundance, in the unburned control. Diversity of plant growth forms in the plots was lowest in the low intensity burned plot but was similar among the others (Table 1).

**Table 1.** Abundance and diversity of growth forms in plots of different fire intensities, 5½ years after the fire. Abundance is shown as frequency in 100 1-m^2^ subdivisions of each plot.

| Growth Forms | Fire Intensity |  |  |  | Control |
| --- | --- | --- | --- | --- | --- |
|  | Very High | High | Medium | Low |  |
| Tussocks | 75 | 99 | 100 | 100 | 79 |
| Upright Shrubs | 76 | 15 | 20 | 10 | 73 |
| Cushion/Mat | 87 | 12 | 39 | 5 | 2 |
| Prostrate Herbs | 81 | 40 | 34 | 12 | 51 |
| Erect Herbs | 16 | 9 | 26 | 5 | 21 |
| Giant Basal Rosette | 51 | 26 | 32 | 8 | 42 |
| Giant Stem Rosette | 74 | 74 | 82 | 61 | 91 |
| Prostrate Shrubs | 33 | 29 | 44 | 6 | 34 |
| Acaulescent Rosette | 1 | 1 | 7 | 3 | 0 |
| Trailing Herbs | 0 | 0 | 0 | 0 | 2 |
| Shannon Diversity | 2.00 | 1.81 | 1.99 | 1.49 | 1.90 |
| Simpson-Gini Diversity | 0.86 | 0.80 | 0.84 | 0.68 | 0.84 |

In terms of overall growth form composition, the plots were 69–90% similar but the low intensity plot was the most distinct from the others, particularly the very high fire intensity plot (Fig. 3). However, there was no clear trend with respect to the fire intensity gradient. *Espeletia* plants which survived the very high intensity fire plot had smaller rosettes and were shorter than in the other plots, though there was no clear trend associated with fire intensity in the rest of the plots (GLM ANOVA for plant height, *F*_4,147_ = 20.42, *p* < 0.0001; Kruskal-Wallis Test for rosette diameter, *X*_4_^2^ = 49.16, *p* < 0.0001; Fig 4).

**Fig. 3.**
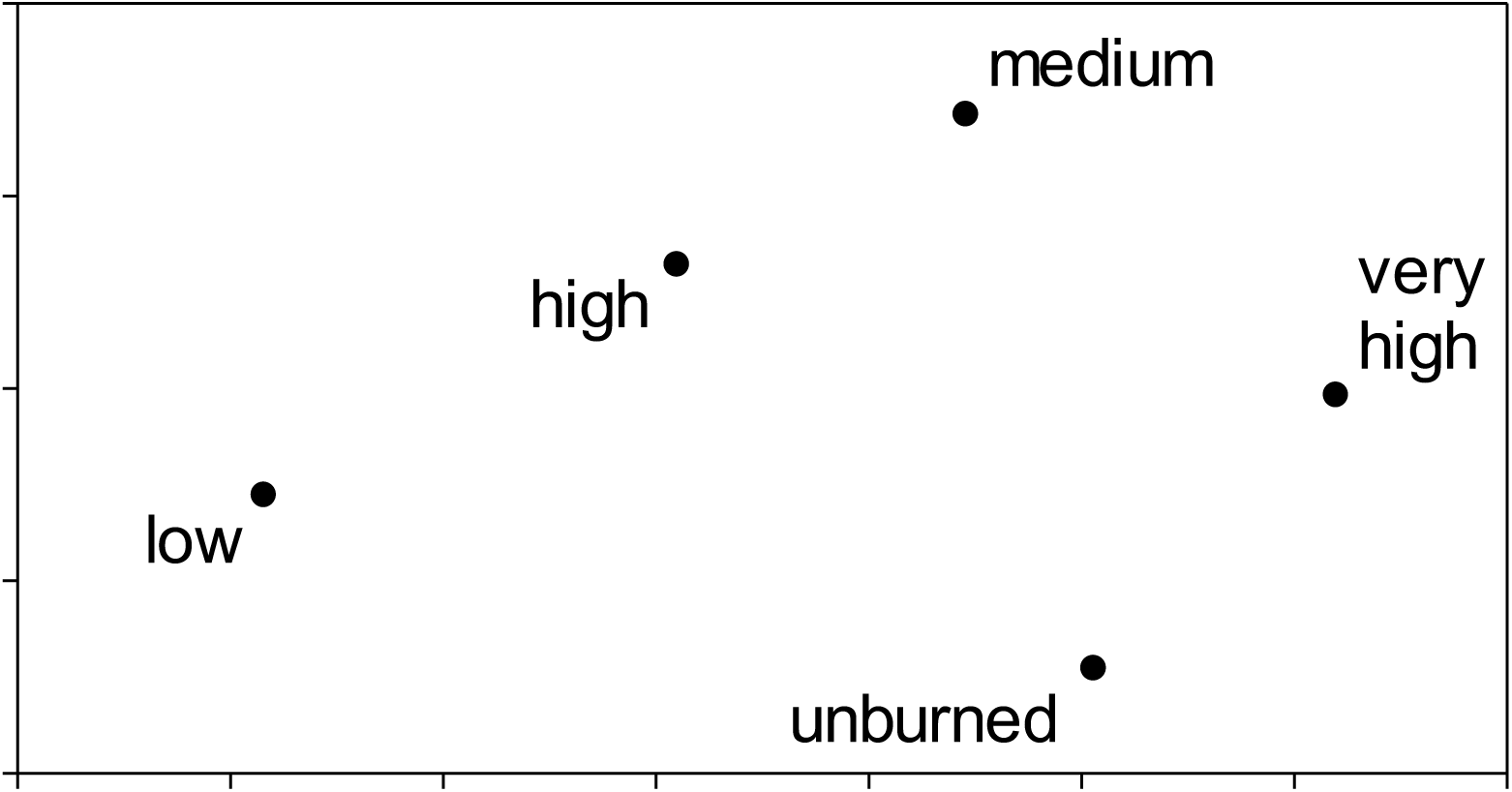
Similarity in plant growth form composition of plots of different fire intensities, 5½ years after the fire. Compositional similarity is represented as distance using non-metric multidimensional scaling: points closer together are more similar in composition. Stress was <0.01.

**Fig. 4.**
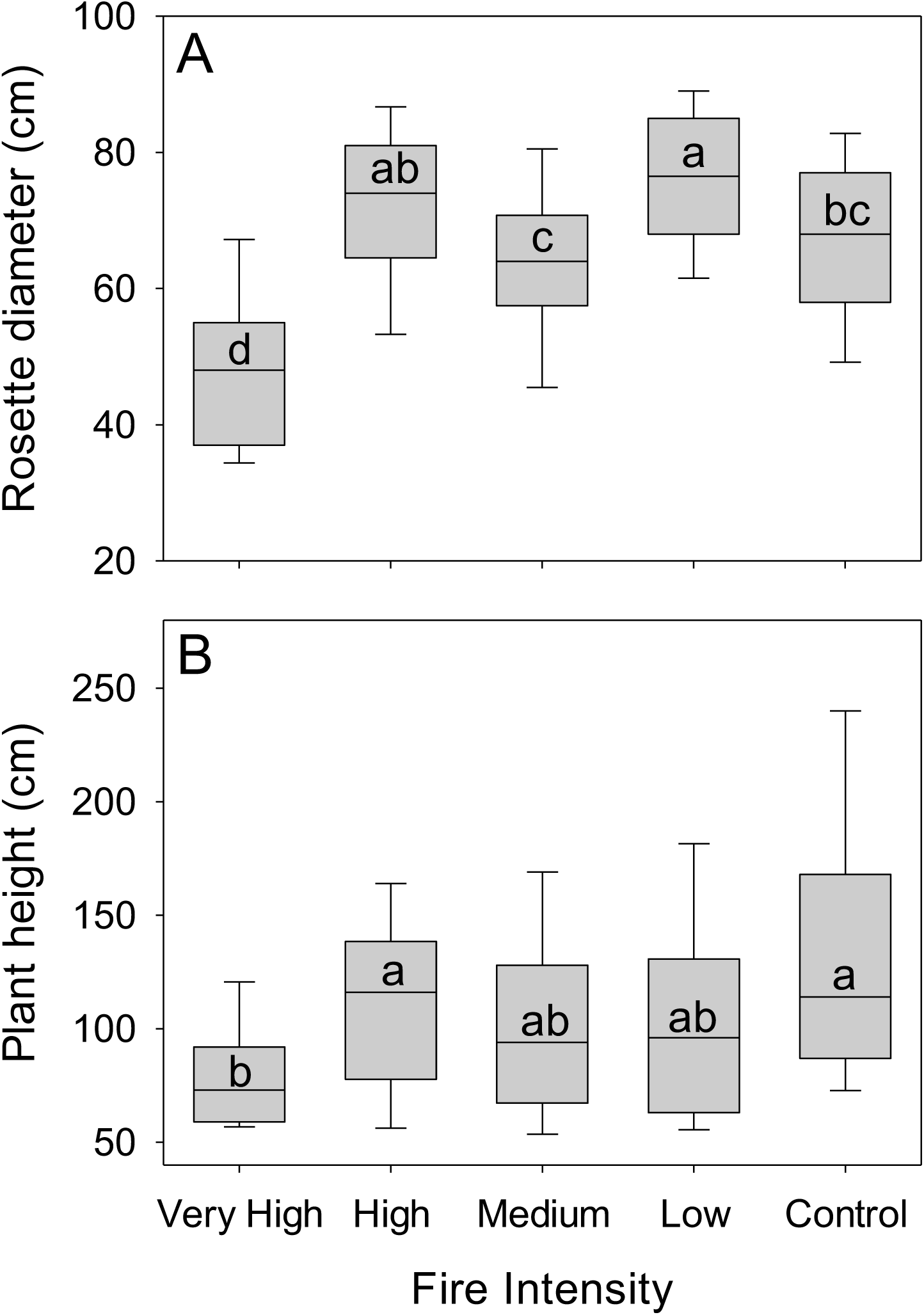
Rosette diameter (A) and height (B) of surviving *Espeletia* plants in plots of different fire intensities, 5½ years after the fire. The control plot had not been burned for at least 40 y. Only plants >50 cm tall at the time of survey were included, since shorter plants were assumed to have arrived after the fire event. Representations of percentiles are as described for Fig. 2. Medians sharing a letter in each panel were not significantly different (*p*≤0.05).

The number of plants which had survived the fire (plants >0.5 m tall at the time of survey) did not show a clear relationship with fire intensity (Fig. 5, white), but there was a clear mortality effect in the very high intensity plot compared with the low intensity plot, with the other plots intermediate (Fig. 5, black). Recruitment of *Espeletia* plants after the fire was lowest in the very high and low intensity plots, and higher in the plots of intermediate fire intensity (Fig. 5, grey). The unburned control plot had a similar number of new recruits to those found in the highest-recruiting burned plots.

**Fig. 5.**
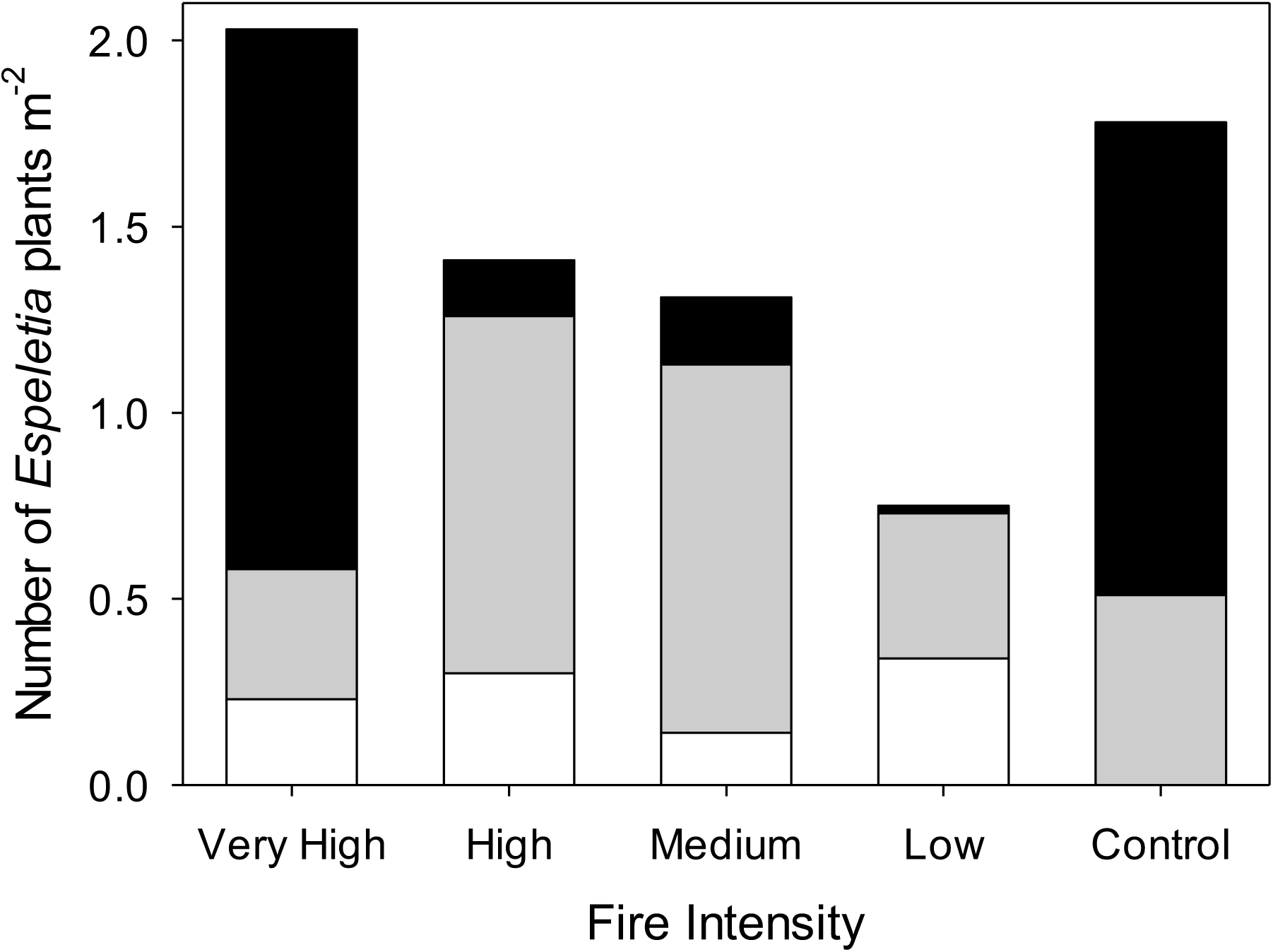
Number of *Espeletia* plants m^-2^ in plots of different fire intensities, 5½ years after the fire. White = plants assumed to have been present during the last fire (plants >50 cm tall at the time of survey); grey = plants assumed to have colonised after the last fire; black = standing dead plants.

## Discussion

The impact of fire intensity on páramo vegetation was still evident 5½ years after the fire: clear differences between very high and low intensity plots in terms of micro-environmental measurements, which were reflected in the growth form composition of the vegetation and in *Espeletia* population structure.

Tussocks showed little variation in response to fire intensities except at the very high intensity, where their abundance was much lower. Tussock grasses are typically the most competitive growth form in páramo plant communities and are usually able to re-sprout after fire (Ramsay and Oxley 1996). It seems that only very high intensity fires generate lethal temperatures in the tussock bases, killing parts or whole tussocks, and thereby stunting tussock grass recovery (Ramsay and Oxley 1996). This is consistent with findings by Ghermandi et al. (2013) that showed high severity fires reduced the perennial herb and perennial grass and shrub cover in Patagonian steppe grasslands due to the mortality of entire or partial bud banks of these re-sprouter species.

*Espeletia* maintained relatively high abundances in all fire intensities but was most abundant in the intermediate intensities that allowed for the survival of *Espeletia* adult plants, while still providing adequate space and light for successful establishment of a post-fire germination pulse. Verweij and Kok (1992) also reported *Espeletia* higher recruitment after fire in a Colombian páramo. Lower intensity sites had lower juvenile recruitment, since these fires move faster and leave some patches unburned (Ramsay 2014), and thus remove less of the tussock canopy. The greatest abundance of juveniles, however, was found in the intermediate rather than very high intensities. This suggests that while fire is an important disturbance factor to create germination space, there is a threshold of fire intensity after which recruitment is no longer enhanced. This might be explained by heat penetration into the soil and consequent seed mortality (Bond and van Wilgen 1996; Fidelis et al. 2010; Horn 1997).

Very high intensity fire also resulted in shorter plants and smaller rosettes of *Espeletia*. Higher mortality rates of taller *Espeletia* plants immediately after the fire in this plot (Ramsay 2014) explains the shorter population 5½ years later. The mortality of these older plants is likely associated with the gradual loss of dead leaves that protect the stem, either by decades of decomposition nearer the base or by gradual removal in previous fires. In a high intensity fire, younger plants with thicker dead leaf cover are more likely to survive (Laegaard 1992; Ramsay 2014; Smith 1981). Very high intensity fires might stunt the growth of rosettes by causing long-term damage to the meristem or modifying environmental conditions that affects the rosettes growth. In the absence of fire, *Espeletia* plants occasionally show pronounced temporary reductions in rosette diameter, with later recovery, perhaps in response to disease or predator attack (personal observations of the authors). Small-diameter rosettes in adult plants are, therefore, potential signals of plants under stress.

Upright shrubs, giant basal rosettes, cushion/mats, prostrate shrubs, prostrate herbs, and erect herbs all increased in frequency as fire intensity increased. These growth forms take advantage of reduced competition from tussocks, the higher availability of light, and warmer soils. Some upright shrubs re-sprout well after fire, from insulated buds or roots, and are known to thrive in fire prone environments (Horn and Kappelle 2009; Ramsay and Oxley 1996). The frequency of shrubs in our survey was measured by rooted presence, thus our result showed an increase in plant number, rather than increasing canopy size. It is likely that it was driven by a combination of re-sprouting from root systems (with relatively rapid growth) and new recruitment from seed, despite the rapid return of tussock canopy cover after fires (Gutiérrez-Salazar and Ramsay in press).

Adults of *Puya hamata* giant basal rosettes are known to survive fire very well, even in high intensity fires (Laegaard 1992). Garcia-Meneses and Ramsay (2014) surveyed *Puya hamata* survival rates immediately after the fire in the same plots we visited, and found only 4.3% of rosettes died (all of them were small plants) in the very high intensity plot. García-Meneses (2012) demonstrated that *Puya hamata* seeds only germinated when soil temperatures exceeded 13°C (around 3°C above ambient daytime air temperatures in our study plots), so the higher light and warmer soil temperatures after higher intensity fires would be expected to promote germination. In the same way, it is likely that cushion/mats, prostrate shrubs, prostrate herbs, and erect herbs suffered increased mortality with greater fire intensities, but this was outweighed by higher rates of subsequent recruitment due to the more open tussock canopies after higher intensity fires.

Acaulescent rosettes were present in relatively low numbers in the burned plots, and completely absent from the unburned control. It is expected that this growth form is more light demanding, and favours the canopy gaps created by burning (Zomer and Ramsay in review). Trailing herbs were only present in the unburned control and absent from all burned plots. Since they often depend on intact tussock grass canopies to climb, it is possible they need more time to establish than 5 ½ years after fire, but it is also possible that seed dispersal in our study area was low.

It is important to note that our study did not look at detailed composition of plant species. Our growth form approach does not reveal if there is a turnover of species within each growth form between intensities. It is therefore very likely that compositional differences between fire intensity plots would be more pronounced (but also noisier) at the species level.

The unburned plot had the most heterogeneous vegetation structure, light at ground level was relatively high, and upright shrubs were observed to be the dominant growth form rather than tussocks. The populations of *Espeletia* in the unburned area were similar in density to those found in some of the burned plots, with many tall *Espeletia* as well as an abundance of *Espeletia* seedlings. This suggests that without fire disturbance upright shrubs can become codominant with tussocks, reducing the canopy cover of the tussocks, and allowing enough light to filter through for *Espeletia* germination and growth. This is a very interesting result, as it has been previously assumed that *Espeletia* would not thrive without fire (Laegaard 1992), and raises some interesting questions about the successional trajectory of paramo plant communities (see Zomer and Ramsay in review).

Our assessment of páramo fire intensity has demonstrated that the same fire can produce different environmental conditions, plant communities, and population structures in different parts of the same fire event. Real fires consist of a patchwork of different fire intensities, the extremes of which produce very different ecological outcomes from intermediate intensities. Clearly, fire management policies for páramo grasslands should consider fire intensity as well as fire frequency (Armenteras et al. 2020; Bremer et al. 2019). A major obstacle to taking fire intensity into account is that it is difficult to measure. Fire intensity is usually measured during the fire event itself. In a study of African savannas, Govender et al. (2006) estimated fire intensity from measurements of fuel loads, fuel moisture contents, rates of fire spread and heat yields of fuel in experimental plots. However, direct measurement of intensity in the páramo will not be possible in most cases, given the remoteness, difficult terrain, cloudy skies (blocking the use of satellite imagery) and lack of investment in research.

Various surrogate indicators have been used in some grasslands as post-fire measures of fire intensity and severity, including leaf and bark scorch height, skeleton twig diameter, ash deposition and plant mortality (Ghermandi et al. 2013; Govender et al. 2006). In the páramo, *Espeletia* plants can act as reliable indicators of time since fire (Zomer and Ramsay 2018), but our current study has demonstrated that *Espeletia* size and population structure did not act as a reliable indicator of fire intensity, except for very high fire intensity. Light transmission to ground level did show a consistent pattern along the full scale of burned intensity plots (very high, high, medium, low). Light decreased as fire intensity decreased. A promising direction would therefore be to focus on light transmission to ground level as an indicator of post-fire intensity. However, there may be a time limit after which these signals can no longer be picked up. It would be useful to follow up this study by continuing to monitor the fire intensity sites assessed in this study to see how long these patterns persist after fire. A complicating factor is that grazing by livestock follows burning in many páramos, and has been shown to further open up vegetation canopies and reduce *Espeletia* densities (Verweij and Kok 1992; Zomer and Ramsay in review).

Despite the difficulties in determining fire intensity, changes in climate and land use are likely to increase the intensity of páramo fires (Armenteras et al. 2020; Keating 2007). Fire suppression policies could result in high fuel loads and greater fire intensities when fires do happen, promoted by more extreme weather events and higher numbers of tourists in the mountains. Our study has shown this could change vegetation composition and the balance of some microenvironmental conditions. Greater proportions of páramo with reduced tussock cover, more bare ground, and higher soil temperatures could have consequences for ecosystem function. High vegetation cover of tussock grasses is often associated with protecting páramo soils that provide water regulation and store and sequester soil carbon (Bremer et al. 2019; Minaya Maldonado 2017). Molina et al. (2019) have linked tussock cover to chemical weathering of páramo soils. More consideration needs to be focused on the impacts of fire suppression policies in the longer term on fire intensities, and the impacts of fire intensities on ecosystem function and service provision.

Our study was provoked by good fortune—being present during a páramo fire with a wide range of fire intensities—but such luck cannot form the basis for a research programme that is badly needed to inform páramo management. Instead, we require a field programme of experimental fires, covering a wide range of fuel loads and land uses. As an illustration of the value of this approach, we refer readers to work carried out in Australian grasslands (Cruz et al. 2018). Very few experimental burns have been carried out in the páramos so far (Horn and Kappelle 2009; Keating 1998; Ramsay and Oxley 1996), but a well-designed experimental programme could form the basis for detailed modelling of fire behaviour and also provide years of subsequent value for monitoring the responses of biodiversity and ecosystem functions to those fires.

## Acknowledgements

This work was carried out as part of permit MAE-DPAC-UPN-BD-IC-FLO-2015-004, issued by the Ecuadorian Ministry of Environment (Carchi province). Fieldwork was carried out by the authors, with assistance from Anna Masters, Cheryl McAndrew, Patricia Gutierrez Salazar, Alejandro Marchán & Juan Yépez Cardenas. Logistical support in REEA was provided by the reserve’s administration and rangers, who also provided information from fire records.

## Notes

### Competing Interest Statement

The authors have declared no competing interest.

